# The potential of dynamic physiological traits in young tomato plants to predict field-yield performance

**DOI:** 10.1101/2021.03.18.435447

**Authors:** Sanbon Chaka Gosa, Amit Koch, Itamar Shenhar, Joseph Hirschberg, Dani Zamir, Menachem Moshelion

## Abstract

To address the challenge of predicting tomato yields in the field, we used whole-plant functional phenotyping to evaluate water relations under well-irrigated and drought conditions. The genotypes tested are known to exhibit variability in their yields in wet and dry fields. The examined lines included two lines with recessive mutations that affect carotenoid biosynthesis, zeta *z*^*2083*^ and tangerine *t*^*3406*^, both isogenic to the processing tomato variety M82. The two mutant lines were reciprocally grafted onto M82, and multiple physiological characteristics were measured continuously, before, during and after drought treatment in the greenhouse. A comparative analysis of greenhouse and field yields showed that the whole-canopy stomatal conductance (g_sc_) in the morning and cumulative transpiration (CT) were strongly correlated with field measurements of total yield (TY: *r*^2^ = 0.9 and 0.77, respectively) and plant vegetative weight (PW: *r*^2^ = 0.6 and 0.94, respectively). Furthermore, the minimum CT during drought and the rate of recovery when irrigation was resumed were both found to predict resilience.

## 1. Introduction

Water stress is the main factor limiting crop yields worldwide [1–3]. Despite intense research over the last decades, drought tolerance is still a major threat to plant growth and crop productivity [4]. The water balance-regulation mechanisms in plants are critical for stress responses, productivity, and resilience, as reviewed in [6]. This balance is controlled by combining two regulation mechanisms: leaf hydraulic conductance [7,8] and the transpiration [9,10]. Continuous measurement of the first one is still a challenge, but high-throughput functional physiological phenotyping (FPP) analysis can be used for the second one[6], which should be considered when selecting traits for crop improvement and predicting crop performance under certain environmental conditions. Accurate yield prediction is important for national food security and global food production [11] and it also aids policymaking. From the research and development perspective, yield prediction tools would enable breeders to reduce the time and cost required to select the best parent lines and test new hybrids under different environmental conditions[12,13]. Finally, reliable yield prediction would benefit the growers who are the end-users of newly developed, improved varieties, aiding their crop management and helping them to make wise economic decisions [14]. However, early growth-stage prediction of crop yields is a challenging task, in general, and is even more challenging under water stress. Several yield-prediction models have been developed, some of which consider yield as a function of genotype (G) and environment (E) and treat the interaction between the two (G×E) as a noise [15,16]. Some other models address G×E interactions using multiplicative models [17], factor analytic (FA) models and linear mixed models to cluster environments and genotypes and detect their interactions [18– 20]. A recently developed yield-prediction model, which is based on a deep neural network fed with weather and soil-condition data for 2,247 sites and yield data for 2,267 maize hybrids) was found to accurately predict yields[13]. The developers of that system concluded that environmental factors had a stronger effect on the crop yield than genotype did. Thus, early-season yield prediction may require a large amount of data from the soil-plant-atmosphere continuum (SPAC). Plant physiological traits that are most relevant to productivity and are very responsive to environmental conditions are expected to serve as important yield predictors [6].

Recent advances in crop physiology show that under drought conditions, quantitative physiological traits such as stomatal conductance [21], osmotic adjustment, accumulation and remobilization of stem reserves and photosynthetic efficiency are strongly correlated with yield [22–24]. Nevertheless, most of the available models do not include key plant physiological traits, such as g_sc_ and photosynthesis, which contribute to crop productivity [25,26]. These traits are among the primary and most sensitive responses of the plant to the changing environment [27] and this dynamic behavior helps to optimize the plant’s response to changing environmental conditions and probably also helps to maximize yield. For example, the early morning peak in stomatal conductance is proposed as a ‘golden hour’ with the assumption of high CO_2_ absorption while transpiration is low due to the relatively low VPD [6].

Therefore, we hypothesized that having a set of high-resolution and continuous data for many key-physiological traits, measured under different environmental conditions at an early growth stage, could improve our ability to predict the yields of particular genotypes under field conditions. To profile physiological traits that reliably contribute to the yield-prediction model, we used two carotenoid biosynthesis mutants, which affect abscisic acid in roots and revealed yield reduction compared with the isogenic control genotype M82 (see Materials and Methods).

## 2. Materials and Methods

### 2.1 Plant material and the grafting procedure

Tomato cv. M82 seeds [28], the recessive mutant *zeta z*^*2083*^ (ZET) described in[29] and the *tangerine t*^*3406*^ (TAN) mutant described in [30,31] were used. Mutants selected as they displayed stable yield reductions when compared to the M82. Moreover, the xanthophylls violaxanthin and neoxanthin are the precursors for the synthesis of xanthoxin, which is converted to ABA. ABA synthesis in roots has been shown to affect plant growth in various ways. Consequently, the ABA synthesis in roots is compromised. Therefore, as a way of increasing yield variation and evaluation for the relative contribution of root ABA to the phenotypes we measure, we made seven grafting combinations, four hetero grafting in which M82 was reciprocally grafted with ZET and TAN, and three self-grafts for each genotype. These mutant lines have mutations that affect two of the four enzymes reported to convert phytoene into phytoene desaturase (PDS), ζ-carotene desaturase (ZDS), zeta isomerase (ZISO) and carotene isomerase (CRTISO; [32,33].

### 2.2 Open-field experiments

The results presented here are from work that was done in two consecutive growing seasons, 2018 and 2019, at the Western Galilee Experimental Station in Akko, Israel. In those trials, we used a low planting density of one plant per m^2^. In 2018, the experiment involved individual plants in a completely randomized design in blocks, with a minimum of 15 replicates per block. In 2019, the experiment was conducted in plots of 10 plants per 5 m^2^, arranged in a randomized block design. The seedlings were grown at a commercial nursery (Hishtil, Ashkelon, Israel) for 35 days and then transplanted into the field at the beginning of April; wet and dry trials were conducted. Both wet and dry fields started the growing season at field capacity, which represents the maximum amount of water that the soil could hold. In the wet treatment, 320 m^3^ of water was applied per 1000 m^2^ of field throughout the growing season, according to the irrigation protocols commonly used in the area. In the limited-irrigation (drought) treatment, we stopped irrigation 3 weeks after planting, so only 30 m^3^ of water was applied per 1000 m^2^ of field. There was no rain during the experimental period, so the drought scenario was managed entirely via irrigation.

### 2.3 Measurements of yield and yield components

The experiments were harvested when nearly 100% ripened. Plant vegetative weight (PW, g m^-2^) was determined by weighing only the vegetative tissue (after harvesting the fruits) without the roots. Total fruit yield (TY, g m^-2^) per plant or plot included both the red and a few green fruits. Mean of 20 red fruits (FW) was estimated from a random sample of 20 fruits per plant or plot. The concentration of total soluble solids (Brix %) was measured using a digital refractometer and a random sample of 10 fruits per plant or 20 fruits per plot. The sugar output per plant or plot was calculated as the product of Brix and TY.

### 2.4 Pigment extraction and analysis

Fresh samples of root and flower tissues (50 to 100 mg) were harvested and immediately frozen in liquid nitrogen. Carotenoids were extracted and quantified according to protocols described by [34].

### 2.5 Greenhouse experiment using the physiological-phenotyping platform

A greenhouse experiment was conducted in parallel with a field experiment from mid-April to mid-May in 2018. The grafted and well-established seedlings were transplanted into 4-L pots filled with potting soil (Bental 11, Tuff Marom Golan, Israel). Plants were grown under semi-controlled greenhouse conditions with naturally fluctuating light (see Fig. 1A). Whole-plant, continuous physiological measurements were taken using a high-throughput, telemetric, gravimetric-based phenotyping system (Plantarry 3.0 system; Plant-DiTech, Israel) in the greenhouse of the I-CORE Center for Functional Phenotyping (http://departments.agri.huji.ac.il/plantscience/icore.phpon), as described in [35].

**Fig. 1.**
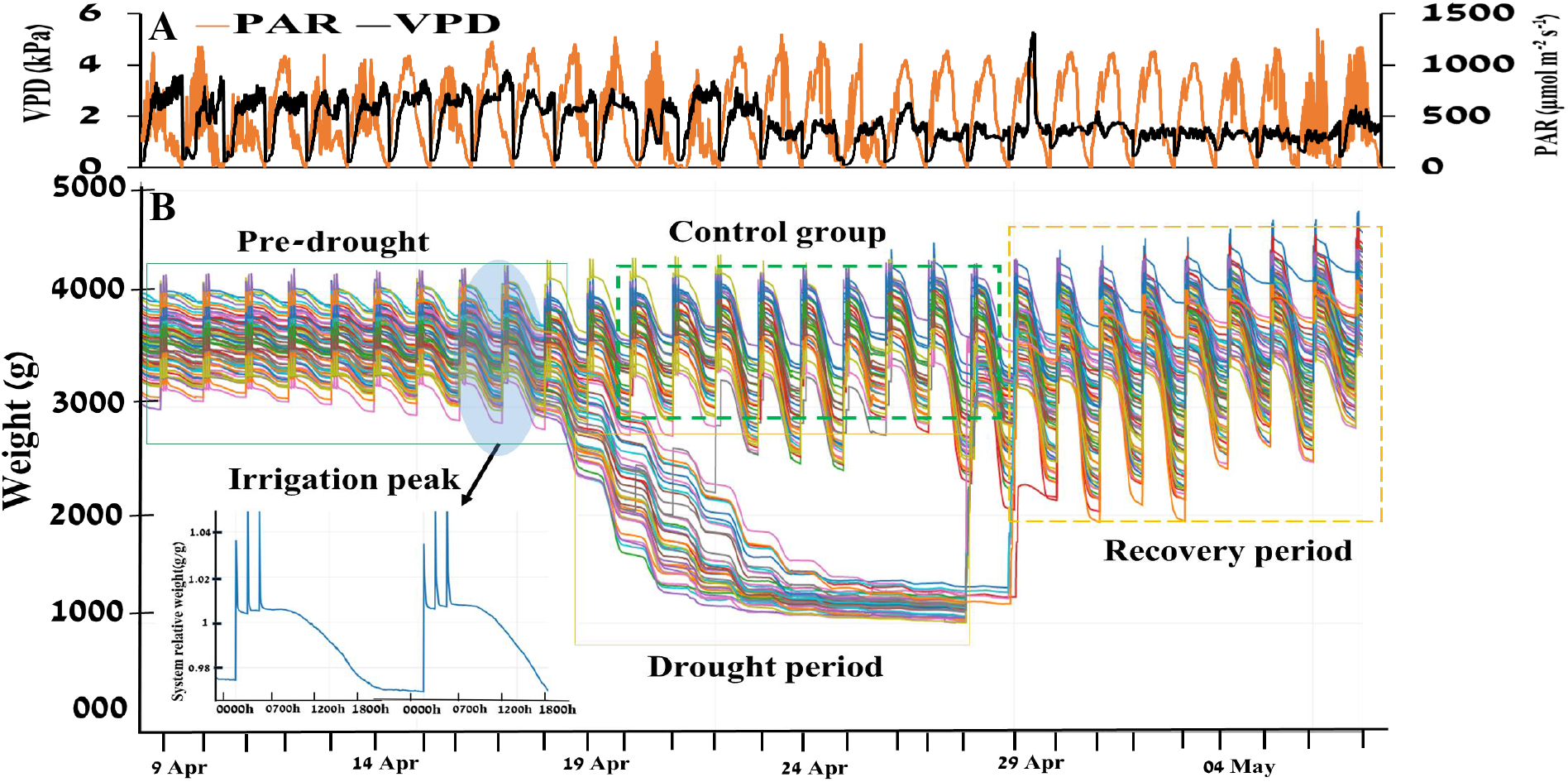
Atmospheric conditions and experimental progress are represented as the fluctuations in pot weight over the course of the experiment in the greenhouse. **(A)** Daily vapor pressure deficit (VPD) and photosynthetically active radiation (PAR) during the 29 consecutive days of the experiment. **(B)** Continuous weight measurements of all the plants during the 29 days of the experiment. Each line represents one plant/pot. The decreasing slope of the lines during the day indicates that the system lost weight as the plants transpired. The three sharp peaks during the nighttime show system weight gain during irrigation events.

The set-up included 72 highly sensitive, temperature-compensated load cells, which were used as weighing lysimeters. Each unit was connected to its own controller, which collected data and controlled the irrigation to each plant separately. A pot containing a single plant was placed on each load cell. (For more details, see the “Experimental set-up” section.) The data were analyzed using SPAC-analytics (Plant-Ditech), a web-based software program that allowed us to view and analyze the real-time data collected from the Plantarray system.

### 2.6 Experimental set-up

The experimental set-up was generally similar to that described by [35], with some modifications. Briefly, before the start of the experiment, all load-cell units were calibrated for accuracy and drift level under constant load weights (1 kg and 5 kg). Each pot was placed into a Plantarray plastic drainage container on a lysimeter. The containers fit the pot size, to enable the accurate return to pot capacity after irrigation and to prevent evaporation. The container had orifices on its side walls that were located at different heights, to allow for different water levels after the drainage of excess water following irrigation. Evaporation from the soil surface was prevented by a cover with a circle cut out at its center through which the plant grew.

Each pot was irrigated with a multi-outlet dripper assembly that was pushed into the soil to ensure that the medium in the pot was uniformly wetted at the end of the free-drainage period following each irrigation event. Irrigation events were programmed to take place during the night in three consecutive pulses (see inset in Fig.1B). The amount of water left in the drainage containers underneath the pots at the end of the irrigation events was intended to provide water to the well-irrigated plants beyond the water volume at pot capacity. The associated monotonic weight loss over the course of the daytime hours was essential for the calculation of the different physiological traits using the data-analysis algorithms (see inset in Fig. 1B).

#### Drought treatment

As each individual plant has a unique transpiration rate based on its genetic characteristics and location in the greenhouse, stopping the irrigation of all plants at once would lead to non-homogeneous drought conditions. To enable a standard drought treatment (i.e., similar drying rate for all pots), drought scenarios were automatically controlled via the system’s feedback-irrigation controller, in which each plant was subjected to a constant reduction in soil water content based on its daily water loss

### 2.7 Measurement of quantitative physiological traits

The following water-relations kinetics and quantitative physiological traits of the plants were determined simultaneously, following protocols and equations [1] implemented in the SPAC-analytics software for daily transpiration, transpiration rate, normalized transpiration (E) and WUE. Cumulative transpiration (CT) was calculated as the sum of daily transpiration for all 29 days of the experiment for each plant. The other physiological traits used in this experiment are described in [36]. The estimated plant weight at the beginning of the experiment was calculated as the difference between the total system weight and the sum of the tare weight of the pot + the drainage container, the weight of the soil at pot capacity and the weight of the water in the drainage container at the end of the free drainage. The plant weight at the end of a growth period (calculated plant weight) was calculated as the sum of the initial plant weight and the product of the multiplication of the cumulative transpiration during the period by the WUE. The latter, determined as the ratio between the daily weight gain and the transpiration during that day, was calculated automatically each day by the SPAC-analytics software. The plant’s recovery from drought was described by the rate at which the plant gained weight following the resumption of irrigation (recovery stage).

### 2.8 Data presentation and statistical analysis

We used the JMP® ver. 14 statistical packages (SAS Institute, Cary, NC, USA) for our statistical analyses. Levene’s test was used to examine the homogeneity of variance among the treatments. Differences between the genotypes were examined using Tukey HSD. Each analysis involved a set significance level of *P* ≤ 0.05.

Pairwise Pearson correlations between traits under greenhouse conditions and the yield and yield components measured in the open field (i.e., plant vegetative weight, red yield, green yield, Brix yield and total yield) were calculated using the genotype’s mean performance.

## 3. Results

### 3.1 Field-based plant weight and total yield

The yield components plant vegetative weight (PW), total yield (TY), and green yield (GY) were tested under well-irrigated and dry conditions in the 2018 and 2019 growing seasons. Comparing two key traits TY and PW we found similar performances of the genotypes across years in 2018 and 2019.

#### Plant vegetative weight (PW)

In the well-irrigated field, the M82 self-grafted plants (M82_scion/M82_rootstock) had a significantly higher PW than the TAN/TAN and ZET/ZET plants. Under the dry condition, no significant difference was observed between the M82 and TAN self-grafted plants, whereas the plant vegetative weights of the ZET/ZET plants were significantly lower (Fig. 2A, B) under both well-irrigated and dry conditions.

**Fig. 2.**
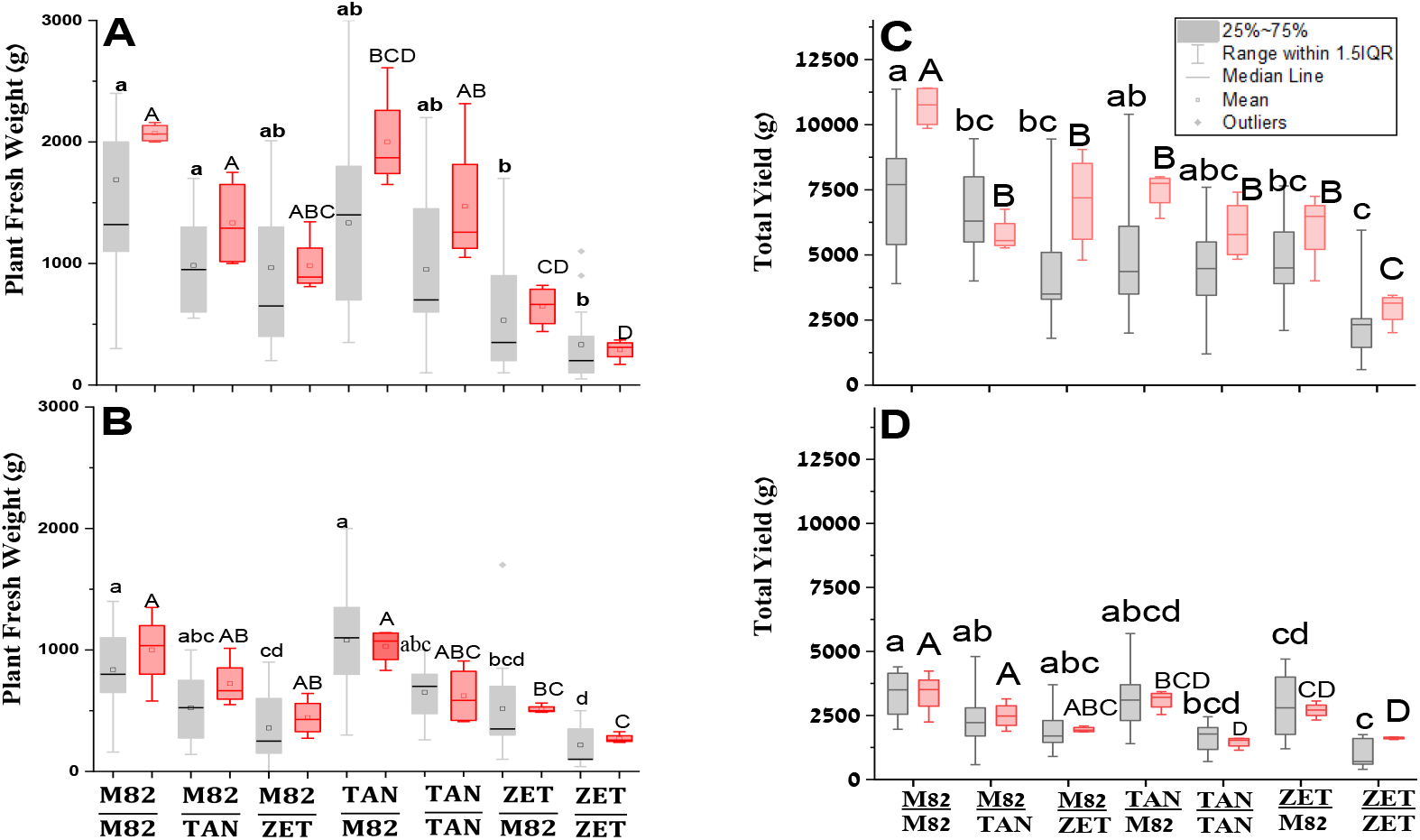
Plant weight and total yield among three reciprocal-grafted tomatoes grown in the field. Boxplot showing the differences in **(A)** fresh weights of self-grafted and reciprocal-graft plants under the well-irrigated condition and **(B)** fresh weights of self-grafted and reciprocal-graft plants under the limited-irrigation condition. **(C)** Total fruit yield self and reciprocal-graft plants under the well-irrigated condition and (**D)** Total fruit yield of self and reciprocal-graft plants under the limited-irrigation condition. Data from 2018 are indicated in grey (with small letters) and data from the 2019 experiments are indicated in red (with capital letters). Different letters indicate significantly different means, according to Tukey’s Honest Significant Difference test (*p* < 0.05). Box edges represent the upper and lower quantile with the median value shown as a bold line and mean as a small square in the middle of the box. Whiskers represent 1.5 times the quantile of the data.s

#### Total Yield (TY)

Under well-irrigated conditions, the TY of the different self-grafted M82 was significantly different from both mutants across both years. The total yield of M82/M82 was significantly higher than the yields of the other self-grafted plants, TAN/TAN was a medium yielder and ZET/ZET had the lowest yield of all the self-grafted plants across both years. Under the drought condition, the total yield of M82/M82 remained higher than those of the other two genotypes, which were not different from each other (Fig. 2C and D, respectively). However, the TY under the drought condition was less than half of that observed under the well-irrigated condition. To increase the phenotypic variation in yield, we used a reciprocal-grafting approach, in which seven combinations of the three tomato cultivars resulted in different gradients of yield performance under wet and dry conditions (Fig. 2C and D, respectively). TY increased more than 2-fold when TAN and ZET scions were grafted onto M82 rootstock, especially under dry conditions.

### 3.2 Early-stage physiological traits measured in the greenhouse

To identify physiological traits of young tomato plants that might serve as good predictors of yield in the field, we profiled multiple physiological traits using continuous data collected on a minute time-scale, such as whole-canopy stomatal conductance (g_sc_); continuous data collected on a daily time-scale, such as transpiration throughout the experimental period as a cumulative transpiration (CT); and single-point measurements such as growth rate and plant net weight (see Table 1).

**Table 1.** Correlations between the physiological traits of young tomato plants in the greenhouse and their field-based yield and biomass under well-irrigated conditions; means of each genotype were used for the correlation. The greenhouse measurements were categorized as continuous (i.e., whole-canopy stomatal conductance, g_sc_,), cumulative or single-point measurements. g_sc_ at the three-time periods (morning, midday, and late afternoon) is obtained by averaging the 3 minutes measurement during each time. All measurements were taken 1 week before the stress treatment started. r^2^ and *p*-values indicate the range of weak to strong correlations.

|  |  |  | Field-Based Measurements |  |  |  |
| --- | --- | --- | --- | --- | --- | --- |
|  |  |  | Total Yield |  | Plant fresh weight |  |
|  |  |  | R<sup>2</sup> range | p-Value ranges | R<sup>2</sup> range | p-Value range |
| Greenhouse-Based Measurements | Continuous | $g_{sc}(7:00-10:00)$ | 0.55 to 0.90 | 0.19 to 0.004 | 0.2 to 0.60 | 0.62 to 0.14 |
| | | $g_{sc}(10:00-13:00)$ | 0.45 to 0.72 | 0.304 to 0.067 | 0.44 to 0.89 | 0.32 to 0.006 |
| | | $g_{sc}(13:00-16:00)$ | 0.34 to 0.71 | 0.447 to 0.073 | 0.74 to 0.93 | 0.054 to 0.002 |
|  | Single point | Cumulative transpiration | 0.77 | 0.04 | 0.94 | 0.001 |
|  |  | Growth rate | 0.62 | 0.12 | 0.89 | 0.0065 |
|  |  | Plant net weight | 0.70 | 0.076 | 0.79 | 0.038 |

The continuous measurement data show that the traits varied with the environment. For example, as shown in Figure 3, the whole-canopy conductance measured every 3 min for the whole day fluctuated over the course of the day in response to the environment. To better understand this trait, we divided the day into three periods: morning, midday, and late afternoon time period. We found that stomatal conductance was relatively high at morning time (Fig. 3, marked in green), declined between midday and late afternoon to some point, and then increased again during the late afternoon. We also performed a correlation analysis using the average value of morning (7:00am -10:00am), (midday, 10:00am-13:00pm) and late afternoon (13:00pm-17:00pm) measurement and correlated it with field-based yield and biomass data.

**Fig. 3.**
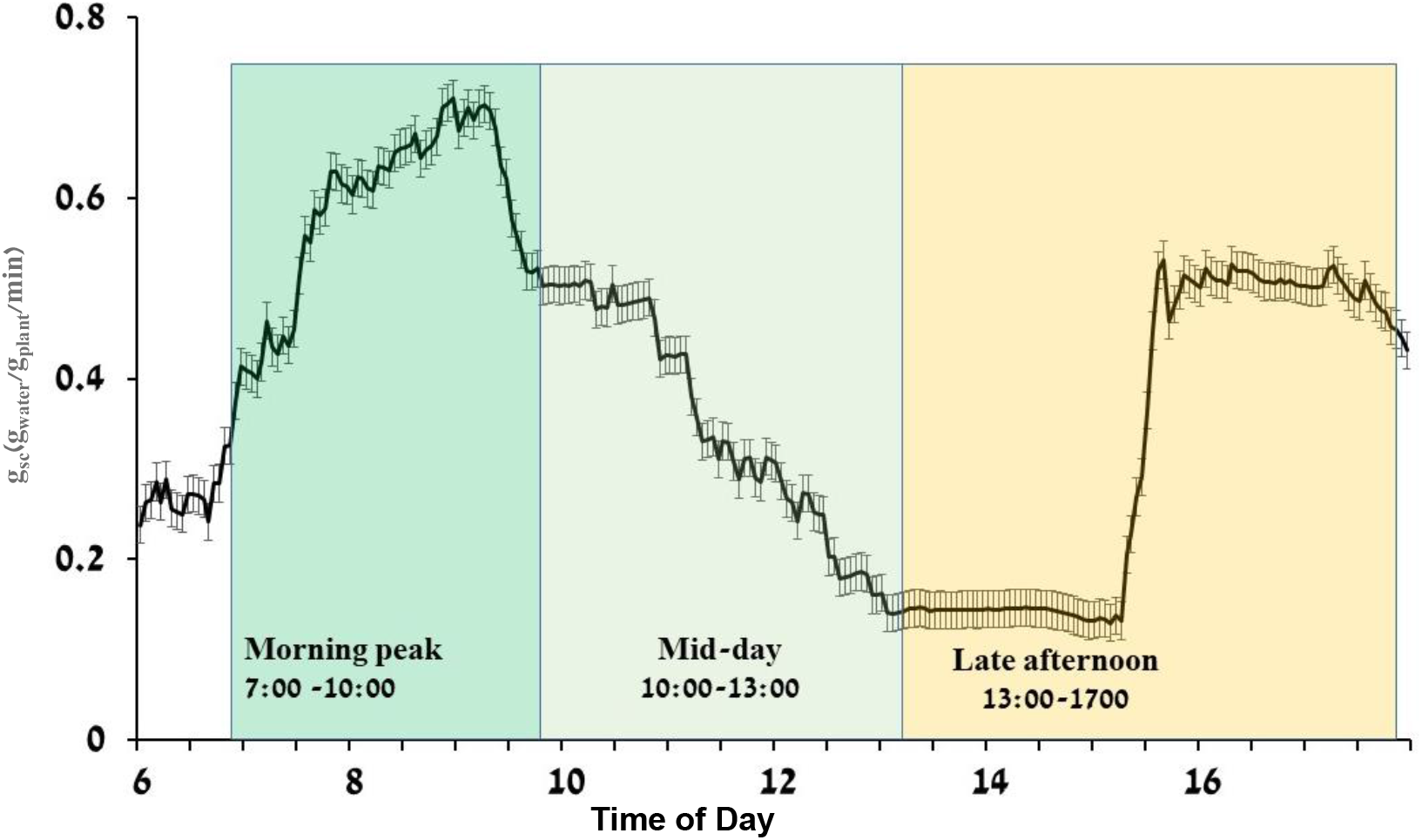
The Daily pattern of whole-canopy stomatal conductance (gsc(g_water_-1g_plant_-1min) presented as an example of continuous whole-plant physiological measurement. Well-irrigated M82 tomato plants were used. The line is an average of three days. Data are shown as means (± SE, n=10).

### 3.3 Correlation of greenhouse physiological traits with yield and yield components in the field

Data from the functional-phenotyping system were composed of continuous soil-plant-atmosphere measurements, with each data point representing the trait at a certain time point. In contrast, field data are normally composed of a single-point measurement that represents the plant’s absolute performance throughout the season (e.g., total fruit yield or plant vegetative weight). When we compared time-series, cumulative and single-point physiological traits (measured traits) of young tomato plants with their field-based yield-related traits (TY, PW, RF, GF and Brix,), we found only a few traits that were highly correlated with each other (Table 1), out of about 95 bivariate combinations (see Fig. 4, Supplementary Figs. S3 and S4). Here, we present a few physiological traits for which the greenhouse data was strongly correlated with the field data and for which we observed low *p*-values (e.g., the highly correlated traits in Table 1).

**Fig. 4.**
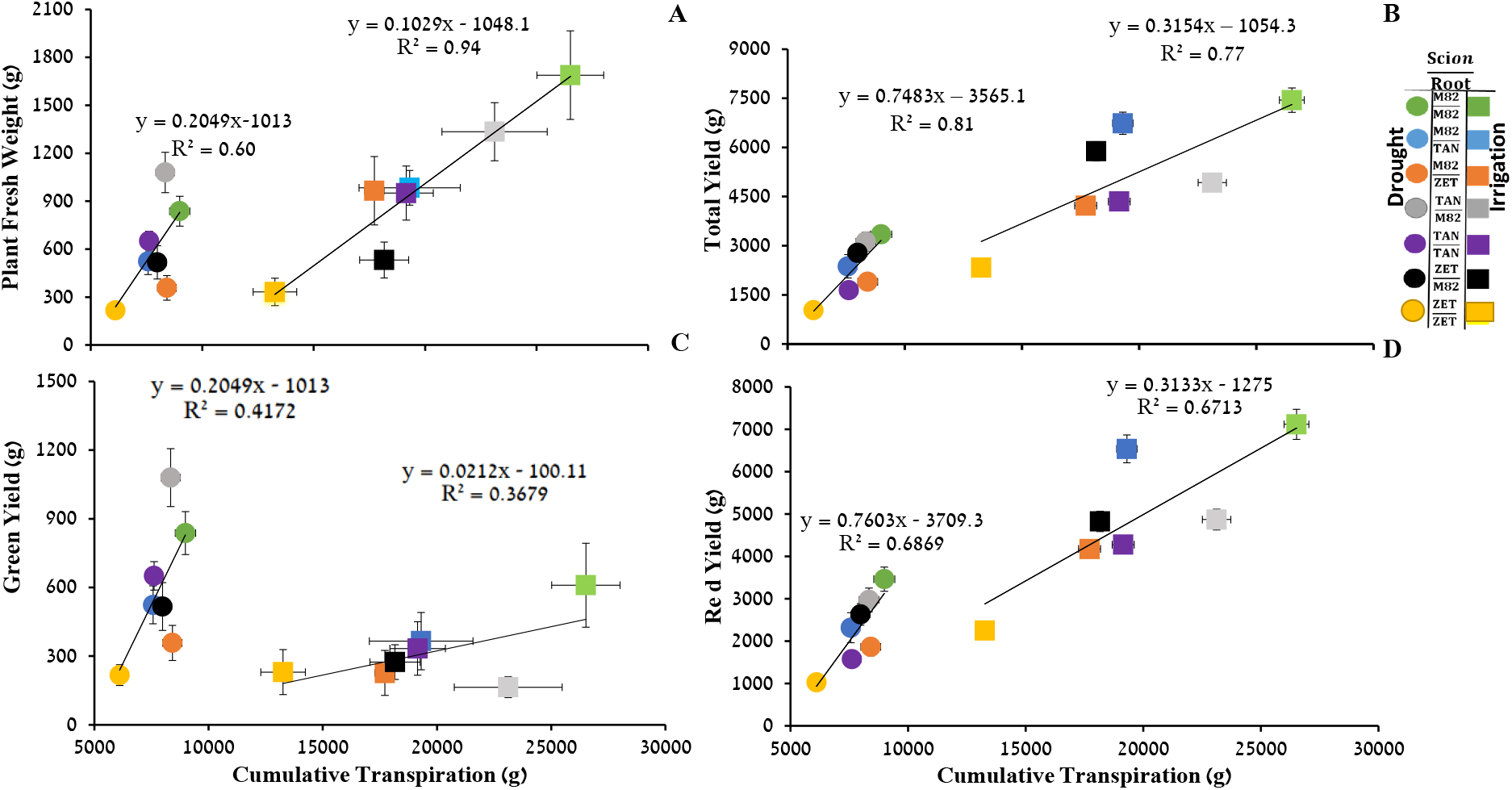
Correlations between yield components and cumulative transpiration of different tomato genotypes. **(A)** Plant vegetative weight in the field, **(B)** total fruit yield, **(C)** green yield and **(D)** red yield. Measurements taken at harvest time were correlated with the CT throughout the 29 days of the greenhouse experiment. Symbols are the means ± SE of traits for each genotype under the limited-irrigation condition (circles) and the well-irrigated condition (square box). Vertical SE (*n* = 12−15), Horizontal SE (*n* = 8−10).

Time-series data are highly dynamic because of the plant’s continuous response to environmental changes (e.g., stomatal conductance, Fig.3; transpiration rate). Therefore, some data points were strongly correlated with yield (e.g., g_sc_ in the morning, Table 1) while others were weakly correlated with yield (e.g., g_sc_ at midday; Table 1). Looking at cumulative physiological data or single-point traits, both presented as a single value (e.g., CT, growth rate, plant net weight), eliminated the need to select a specific time point and revealed highly significant and positive correlations between CT and yield and most of the yield components under well-irrigated conditions (Fig. 4A−D). Similarly, the CT of drought-treated plants after recovery in the greenhouse was positively correlated with yield and with most yield components, but poorly correlated with green yield (Fig. 4C). A similar positive correlation between CT and yield in the field was observed in 2019 (Supplementary Figs. S5 and S6).

### 3.4 Cumulative transpiration as an indicator of resilience and yield performance

The rate of plants’ recovery from drought stress (i.e., drought resilience) is an important trait. To evaluate this resilience, we measured the CT for the first week after recovery from drought. We then compared that CT data with CT data from two other periods during the experiment: the pre-drought period and the drought period (Fig. 5A). While the CT over the pre-drought treatment showed a similar positive correlation with that of the entire well-irrigated experiment (Fig. 5B), we found a strong negative correlation between total yield and CT and under drought conditions (Fig. 5C). We also observed a strong positive correlation between CT and TY during the recovery period (Fig. 5D), even though the actual total yield of the drought-treated plants was half that of the plants grown under the well-irrigated condition.

**Fig. 5.**
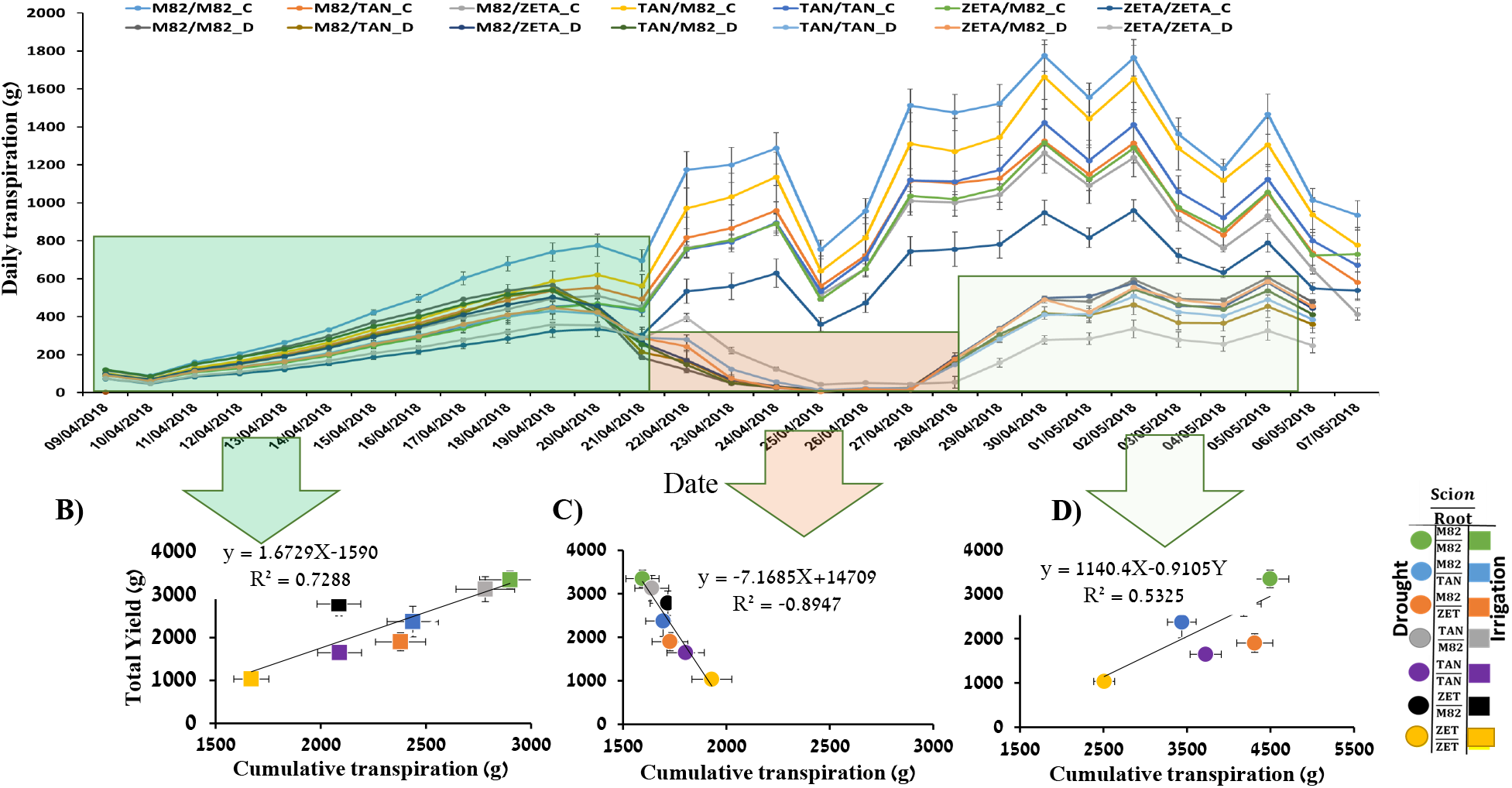
The differential contribution of transpiration periods to yield prediction. **(A)** Mean ± SE. Daily was transpiration continuously measured during the whole experimental period for all genotypes. We examined the relative contributions of the three phases for yield prediction: well-irrigated (green box), drought treatment (orange), and recovery from drought (light green). CT was measured and correlated with TY for each period: **(B)** pre-treatment, **(C)** drought period and **(D)** recovery period. Vertical SE (*n* = 12−15) field-based data; horizontal SE (*n* = 8−10) greenhouse-based data.

## 4. Discussion

Physiological traits (e.g., photosynthesis or stomatal conductance) are key contributors to plant productivity and yield [37,38]. However, existing methods of measuring these traits are mostly manual and thus are limited to a single point on a single leaf at a time [39]. As these physiological traits are very sensitive to ambient conditions, especially light and vapor pressure deficit (VPD); [6], conventional manual measurements fail to capture the temporal and spatial dynamic interactions between the genotype and the environment. This could be misleading for yield prediction, as plants respond differently to dynamic growing conditions [6]. Hence, the integration of manual physiological measurements into breeding programs is limited, most likely due to their low-throughput nature and the large degree of variation within and between temporal and spatial measurements.

In this study, we used continuous measurements of physiological traits to assess whether those traits could serve as early predictors of plant responses to environmental conditions. We used a high-throughput, physiological phenotyping platform, with a high resolution of 3-min intervals, to capture plant responses to the environment. In our experiments, we captured a detailed profile of each plant’s performance. Yet, another challenge was to leverage the daily dynamic responses of plants from these detailed profiles in order to understand their importance in the actual field condition (e.g., choosing the measurement points to be used). A good example of this challenge is demonstrated in Fig. 3, which shows how continuous g_sc_ measurements were correlated with yield performance at different hours of the day (Table 1). Using data from different time periods of the day, we show that the morning g_sc_ peak is strongly correlated with TY and PW in the field. In agreement with our results, a recent study reported high stomatal conductance and photosynthesis in rice in the early morning [40]. The early morning peak has been reported on several plants [9,41] was referred to as a “golden hour” [6], due to relatively low VPD and good light for photosynthesis at this time. In fact, these conditions are allowing the plant to maintain high productivity with low water loss, thereby achieving optimal WUE. As such, we suggest that as soon as the plant reaches this point and as high as its g_sc_ is at this point, it will be more beneficial to the plant in general and in particular under stress. However, a clear understanding of the optimal stomatal conductance kinetics throughout the day and during the entire growing season as it reacts to dynamically changing environmental conditions is still a challenge. Several models have been proposed to understand the kinetics of stomatal conductance at leaf level [42–44] and quite a few at the whole plant level [45]. Although these models are good tools in predicting the kinetics of stomatal response to environment, still it is not easy to leverage the predicted or directly measured small dynamic responses on hourly, daily, and seasonal bases and translate it to final yield. Moreover, the fact that our midday g_sc_ data was less strongly correlated with field performance is in agreement with the common practice of measuring g_sc_ between 10:00 and 14:00 [46,47]. The weak correlation between midday g_sc_ and yield could be related to the dynamic patterns of daily whole-plant water-use efficiency suggested by [6]. Nevertheless, the identification of the best time to measure each trait and/or weather condition understanding the cumulative effect hourly, daily seasonal changes in stomatal conductance on plant performance and dynamic water use might require the use of new data-analysis tools, potentially a data-hungry machine learning algorithms[48], to create a more comprehensive understanding of our large amount of data. However, the application of machine learning in plant science is still in its infancy [48]. Moreover, a better understanding of the genetic mechanism governing the morning peak could contribute to the improvement of crop productivity through breeding, in addition to yield prediction, as plants use water very efficiently at that time of day. It is also important to examine many genotypes. For example, the tomato introgression line (IL) population [28] with multiple years of field data, to verify whether these morning peaks are present in all genotypes, since the current study used only isogenic lines. This would improve our understanding of the genetic mechanism for this important trait.

The relationship between transpiration and net carbon assimilation or dry weight has been well studied [49,50]. The reason for this correlation is most likely due to the fact that CO_2_ enters via the same open stomata through which the plant transpires. Indeed, we found a positive correlation between CT and yield. Yet, this correlation was weaker than the correlation between morning g_sc_ and TY (*r*^2^ = 0.9 versus *r*^2^ = 0.77 and *p* = 0.004 versus *p* =0.04, respectively), suggesting that the correlation between CT and CO_2_ absorption might be affected by other environmental factors, such as VPD, radiation and humidity, which are all known to affect stomatal conductance [51]. On the other hand, CT is a stable, single-point measurement that is relatively simple to measure, yet it integrates the overall responses of plants to the environment throughout the experimental period. Nevertheless, these correlations should be examined in other plant species, as different vegetative stages, reproductive systems, growth, and development patterns may involve different yield-related predictive traits.

Another goal of this study was to evaluate stress-related traits that could predict yield. Under water-deficient conditions, the plant undergoes several changes ranging from molecular and cellular changes to changes at the whole-plant level. This reprogramming of metabolic pathways and physiological response patterns enables the plant to better cope with drought stress [52,53]. Many of the physiological responses to stress [e.g., reduced stomatal conductance, damage to the photosynthetic parts, reduced chlorophyll content; [54]] have dramatic effects on plant productivity. Under stressful conditions, plants enter a protective or survival mode [53] at the expense of their productive mode [55]. Here, we found that CT was strongly and positively correlated with TY under well-irrigated conditions, but negatively correlated with yield under stressful conditions (Fig. 5C). This reversal reflects the productive-survival transition mode of the plant [55]. Namely, a plant that can maximize its transpiration under well-irrigated conditions and swiftly minimize it under stressful conditions is more likely to produce more yield by the end of the season if it recovers quickly after the stress ends. This is clearly shown in Figure 5B: M82/M82 and TAN/M82 had higher levels of transpiration pre-stress, but swiftly reduced their transpiration during the stress period (Fig. 5C) and went back to their high levels of transpiration after recovery (Fig. 5D), which might have led them to have higher yields than the other lines. Thus, this transition mode is important for distinguishing plants’ stress-response (protective) mode from their normal growth response (productive mode). An additional important phase of the plant-stress response is the plant’s post-stress performance, often called resilience.

Resilience to water limitations, specifically the plant’s ability to resume growth and gain yield after water resumption following drought stress, was acknowledged by [56]. Resilience is considered to be a key trait for crop improvement for water stress [57]. Although it has not received much attention for some time [58], this trait has been prioritized in some breeding programs [59]. In this study, we found that the CT of all the treatment periods together (pre-treatment, drought and the recovery period) and the CT of only the recovery period each had a strong, positive relationship with TY (Figs. 4B, 5D), suggesting the importance of this trait for stress-response profiling.

Though the lines TAN and ZET were selected for this study due to their well-characterized yield data, we would like to discuss the contribution of the specific carotenoid mutations to stress response. Carotenoid biosynthesis in roots serve mainly the supply for the abscisic acid (ABA) and strigolactones precursors, β-carotene and violaxanthin, respectively. The mutation *tangerine* (TAN) in the gene *Crtiso* and *Zeta* (ZET) in the gene *Ziso* impair carotene isomerase and ζ-carotene isomerase, respectively. Mutations in these enzymes block carotenoid biosynthesis in their respective states and eliminate downstream xanthophylls in roots (Fig. S2). The accumulation of carotenoid intermediates in roots of TAN and ZET indicates that carotenoid biosynthesis does take place in roots. The low concentration of carotenoids in wild-type roots can be explained, in part, by the synthesis of ABA and strigolactones in the tomato roots, as reviewed in [40]. ABA deficient in roots in mutants TAN and ZET is expected to affect the ability of these plants to cope with drought and other stresses. Our results show that ZET and TAN are prone to slow recovery rates (Fig. 5A, D), in terms of cumulative and daily transpiration, which probably contributes to their low yields. Their lower CT levels may be related to their root systems, since both ZET and TAN grafted as scions on M82 performed a lot better than ZET and TAN when M82 was used as the scion. In these mutants, carotenoid synthesis is blocked, so intermediate products accumulate. This blockage is very effective in the roots due to their lack of exposure to light, whereas exposure to light in the leaves partially compensates for the lack of carotenoid isomerase CRTISO and ZISO [60,61]. However, this cannot explain the lower yields of ZET and TAN under the well-irrigated condition. The relatively low yield of TAN plants under the well-irrigated condition might be linked to the lower concentrations of carotenoids, such as violaxanthin and neoxanthin, in their flowers, as compared to M82 (Supplemental Fig. S2). Violaxanthin and neoxanthin are the precursors for ABA synthesis [62], which suggests that ABA might have been involved in reducing the yields of these mutants. However, this hypothesis needs to be tested in future research.

## 5. Conclusions

In conclusion, continuous measurements of dynamic traits such as g_sc_ provide a dataset that is rich, yet also very challenging to analyze. In our current study, we confirmed that early morning g_sc_ is an important physiological trait that can predict yield performance. Understanding the genetic mechanism underlying early-morning g_sc_ could be a potential avenue for breeding programs aimed at developing lines that will perform well under water-deficit conditions. Furthermore, future data-science tools are likely to improve our understanding of the mechanisms involved and allow us to use these dynamic traits in yield-prediction models. On the other hand, the relatively simple trait of CT of young tomato was proven to be a good predictor of plant biomass and yield performance. The inclusion of CT in yield models is expected to improve the accuracy and consistency of those models, which should facilitate the selection of complex traits for water-stress conditions.

It is important to note that various crops may present different response profiles, as well as different levels of susceptibility to a particular type of stress, depending on their biochemical, physiological and phenological stage.

In addition to yield prediction and crop improvement (i.e., at pre-breeding stages), high-resolution, continuous physiological data could further be exploited to help bridge the genotype-phenotype gap, by combining the functional-genomics approach with a high-resolution time axis on a QTL map. This combined approach may help to identify time-dependent QTLs for dynamic physiological traits such as g_sc_ and help us to understand the genetic mechanisms that underlie those dynamic traits if tested for other crops, since our current work focused only on tomato plants.

## Supplementary figures

**Supplementary Fig. S1.**
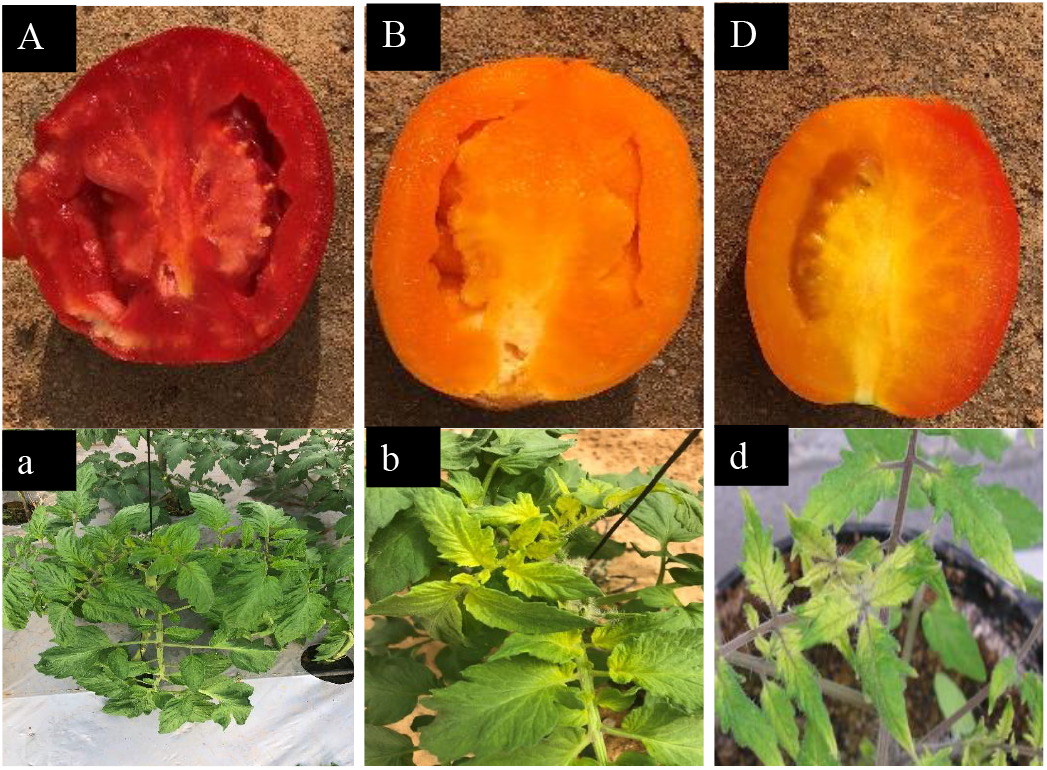
Fruit color and leaf characteristics of M82 and the two mutant lines. **(A, a)** Fruit and leaves of M82. **(B, b)** *Zeta* mutant’s fruit and yellowish leaves characteristic. **(C, c)** The *tangerine* mutant’s fruit and leaf characteristics.

**Supplementary Fig. S2.**
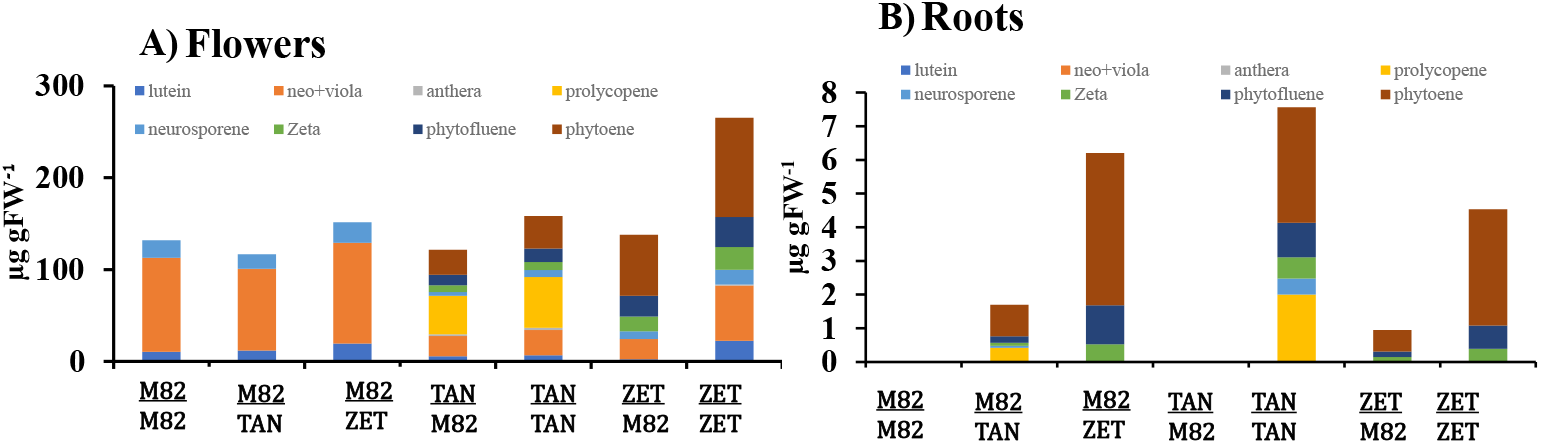
Carotenoid concentrations in the **(A)** flowers and **(B)** roots of wild-type (M82), TAN and ZET plants and their reciprocal combinations (μg. g^-1^ FW). Roots of wild-type tomato contained negligible amounts of carotenoids.

**Supplementary Fig. S3.**
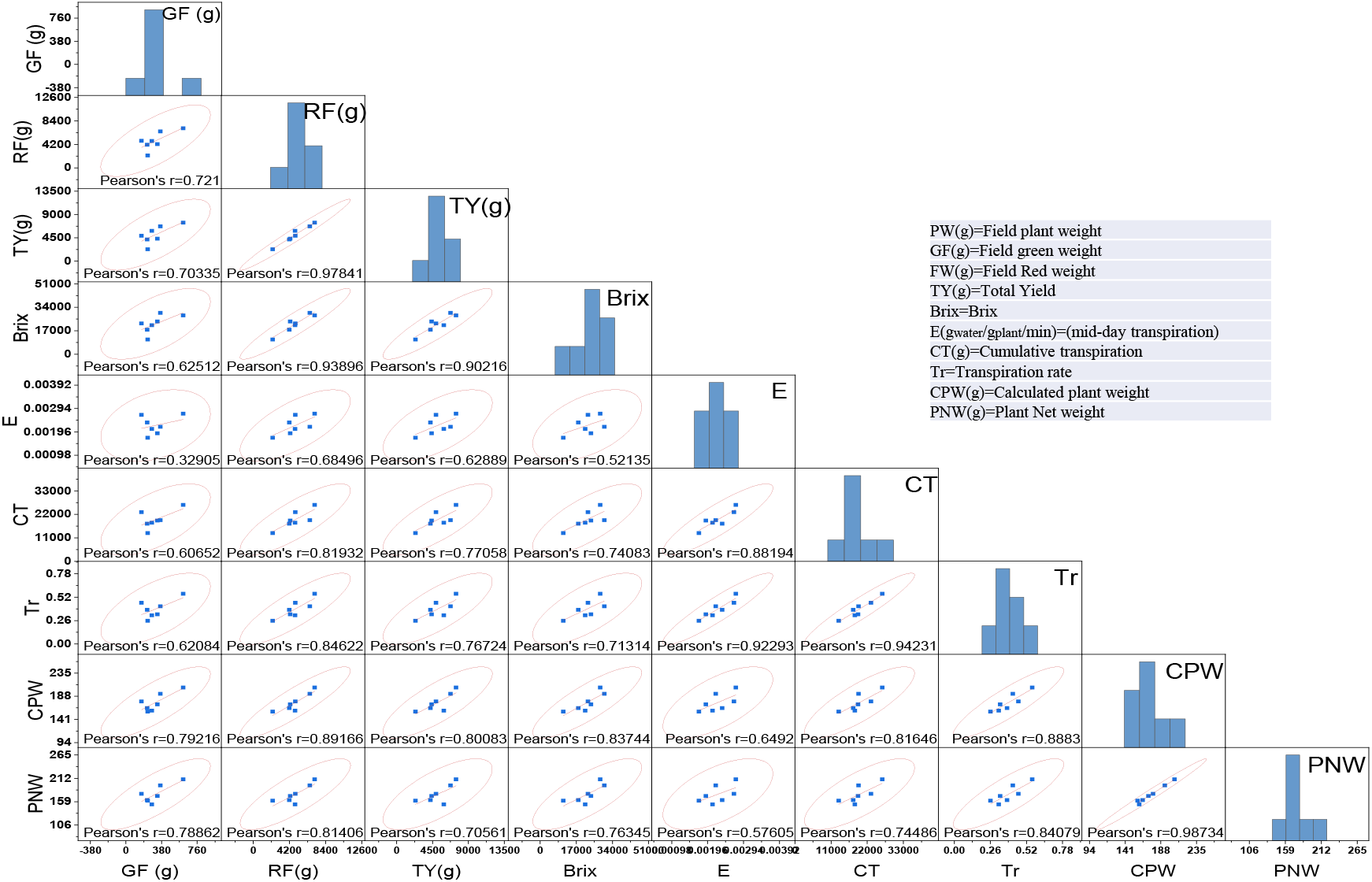
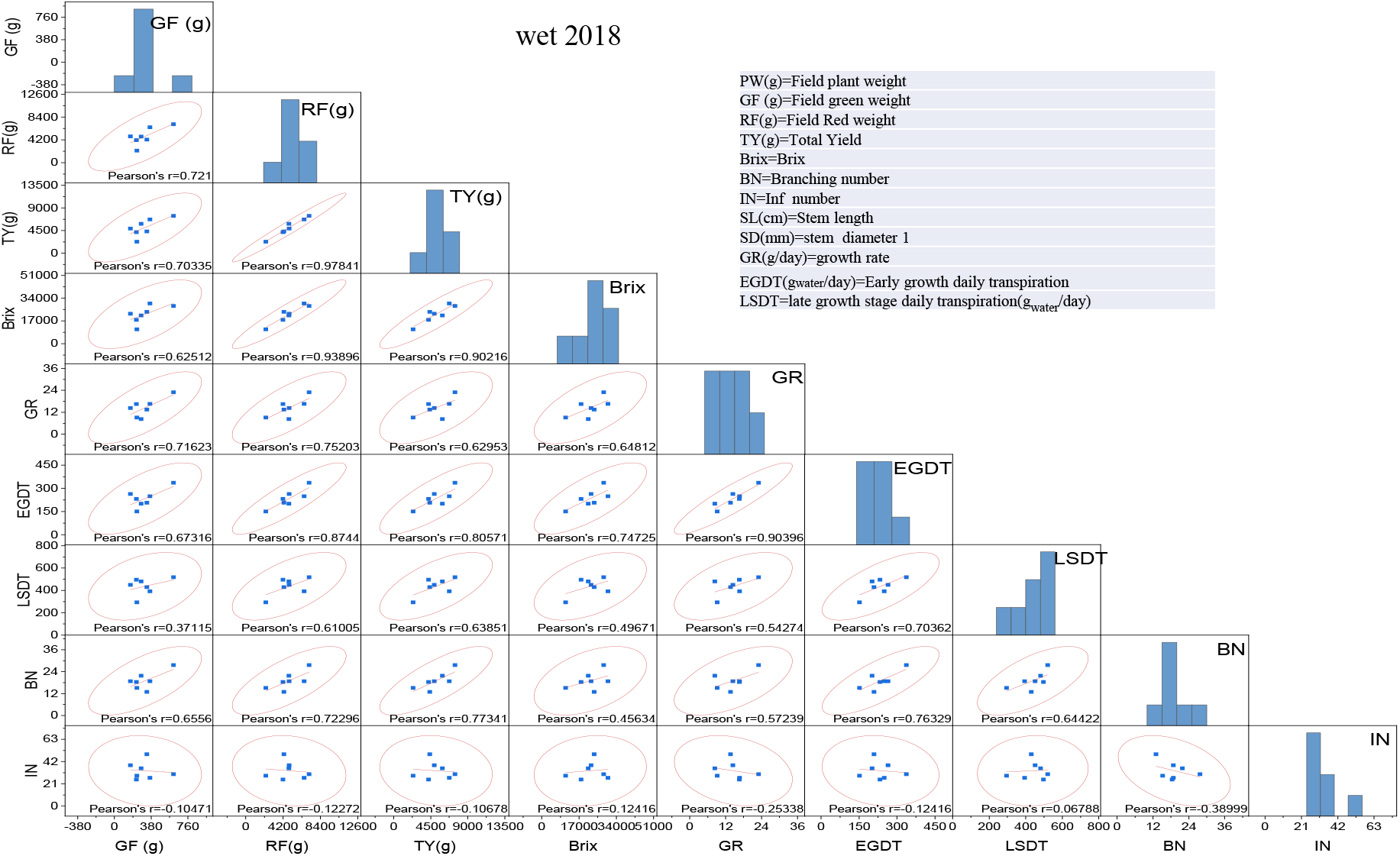
Scatter-plot matrices for traits under the well-irrigated condition in 2018. The figure depicts the matrices of scatter plots and Pearson’s correlation coefficients among the field-measured yield data and yield components correlated with the greenhouse-based traits of 7 different young tomato plants under the well-irrigated condition. The windows show Pearson’s correlation coefficients (*r*) and bivariate scatter plot matrices with a density ellipse. The short names are defined as belo.

**Supplementary Fig. S4.**
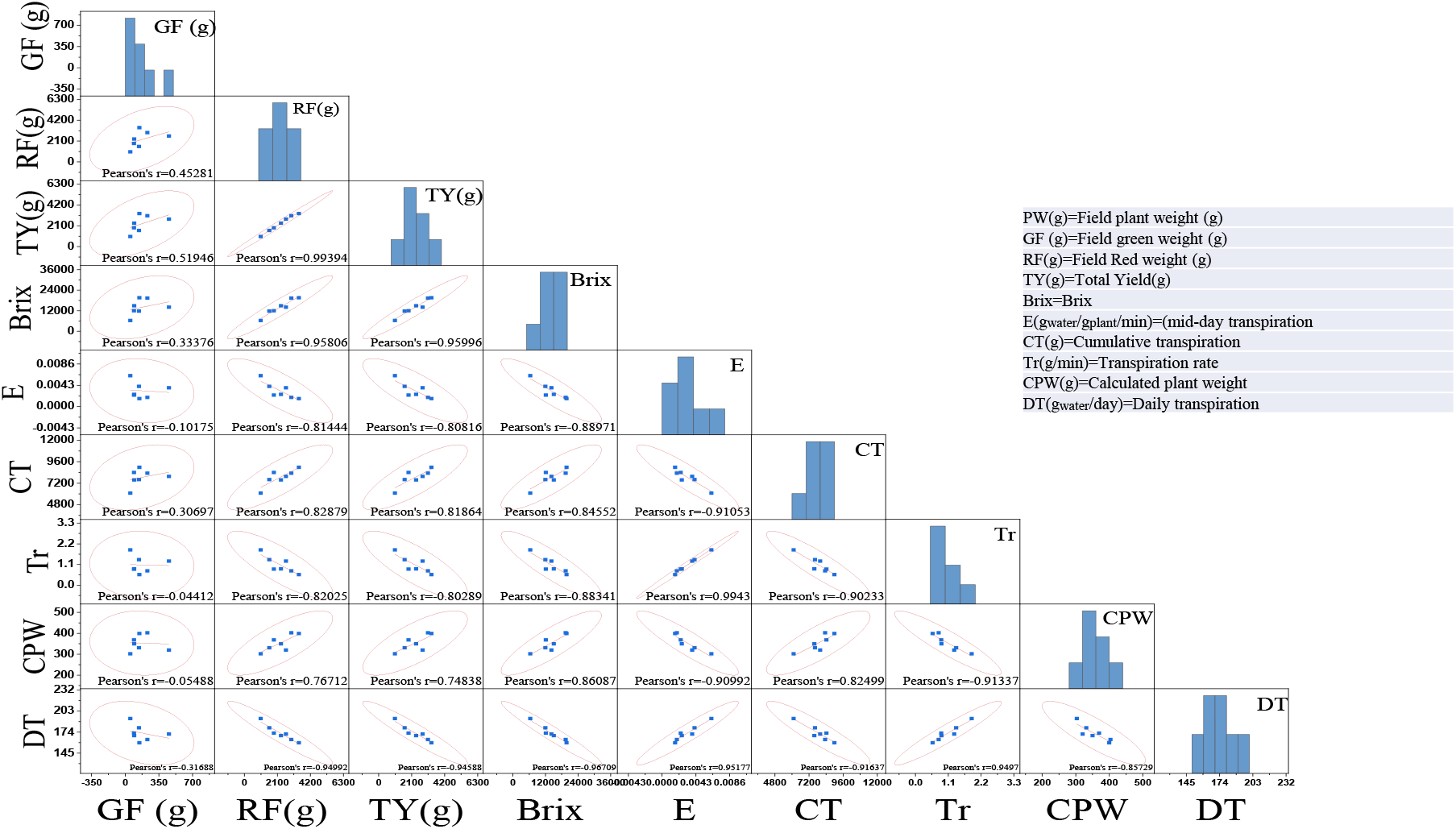
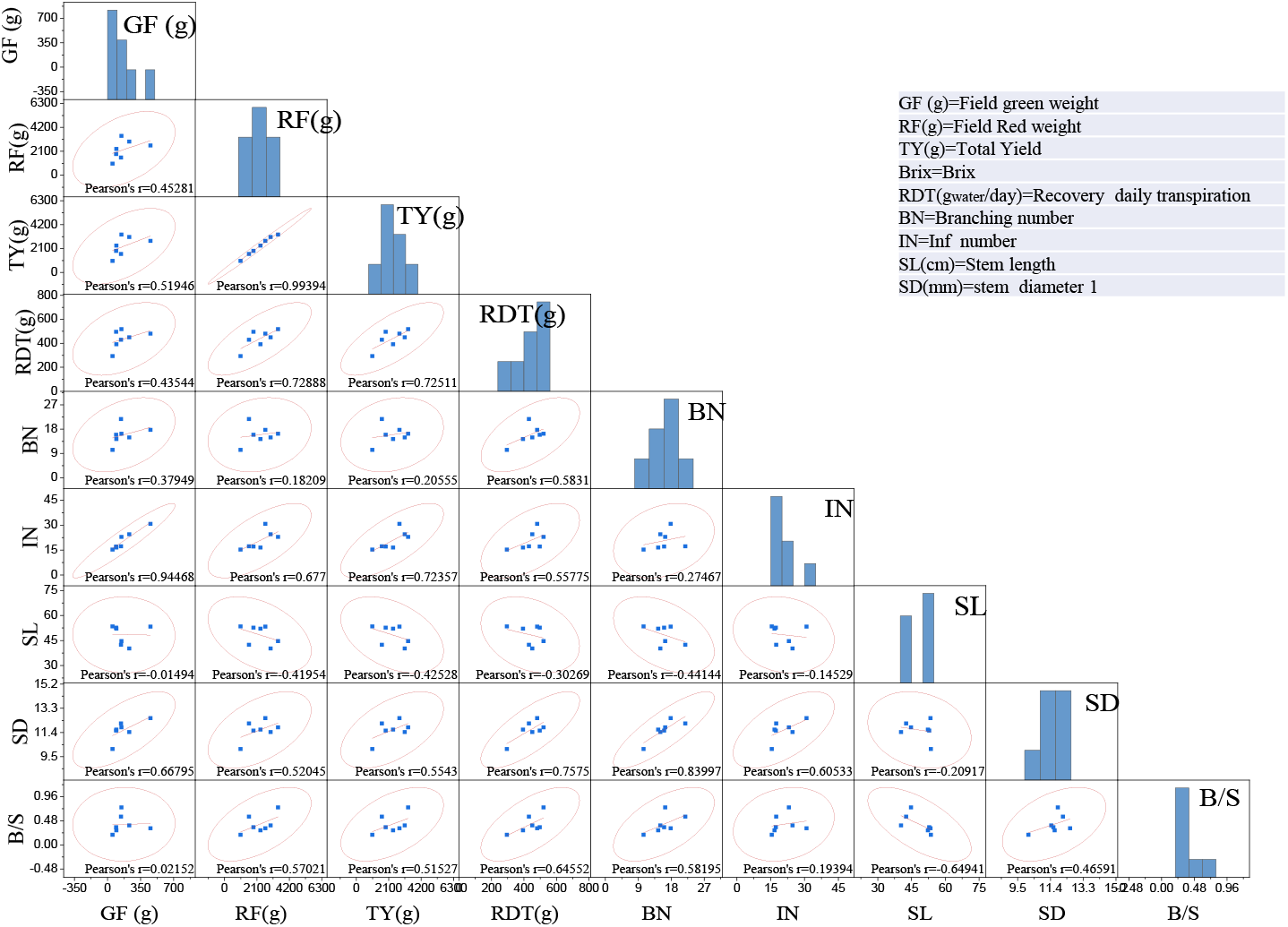
Scatter-plot matrices for traits under the water-deficit condition in 2018. The figure depicts the relationships between field-measured yield and yield components, and the greenhouse-based traits of 7 different young tomato plants that were exposed to drought. The windows show Pearson’s correlation coefficients (*r*) and bivariate scatter-plot matrices with a density ellipse. The short names are defined as below.

**Supplementary Figure 5:**
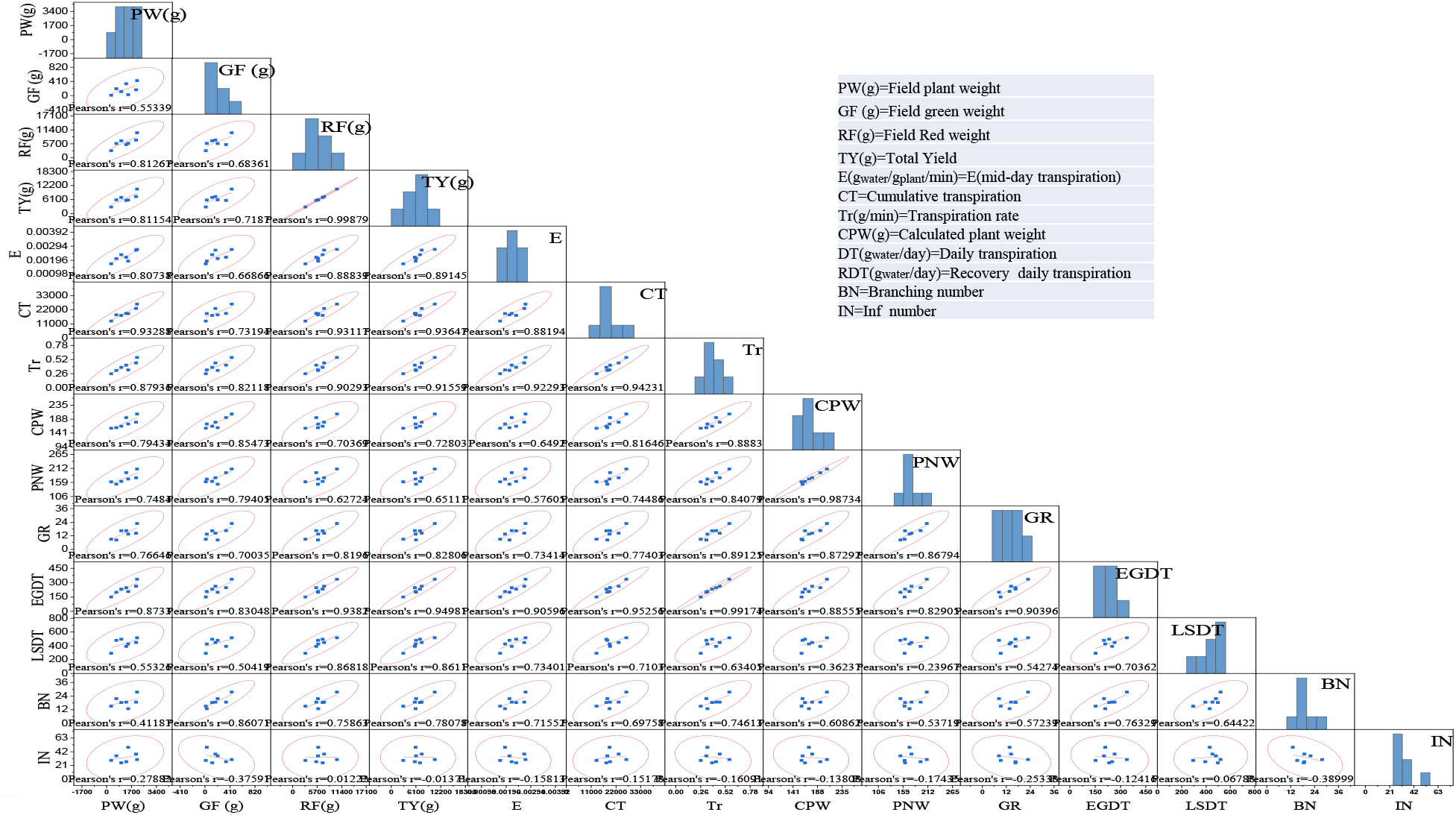
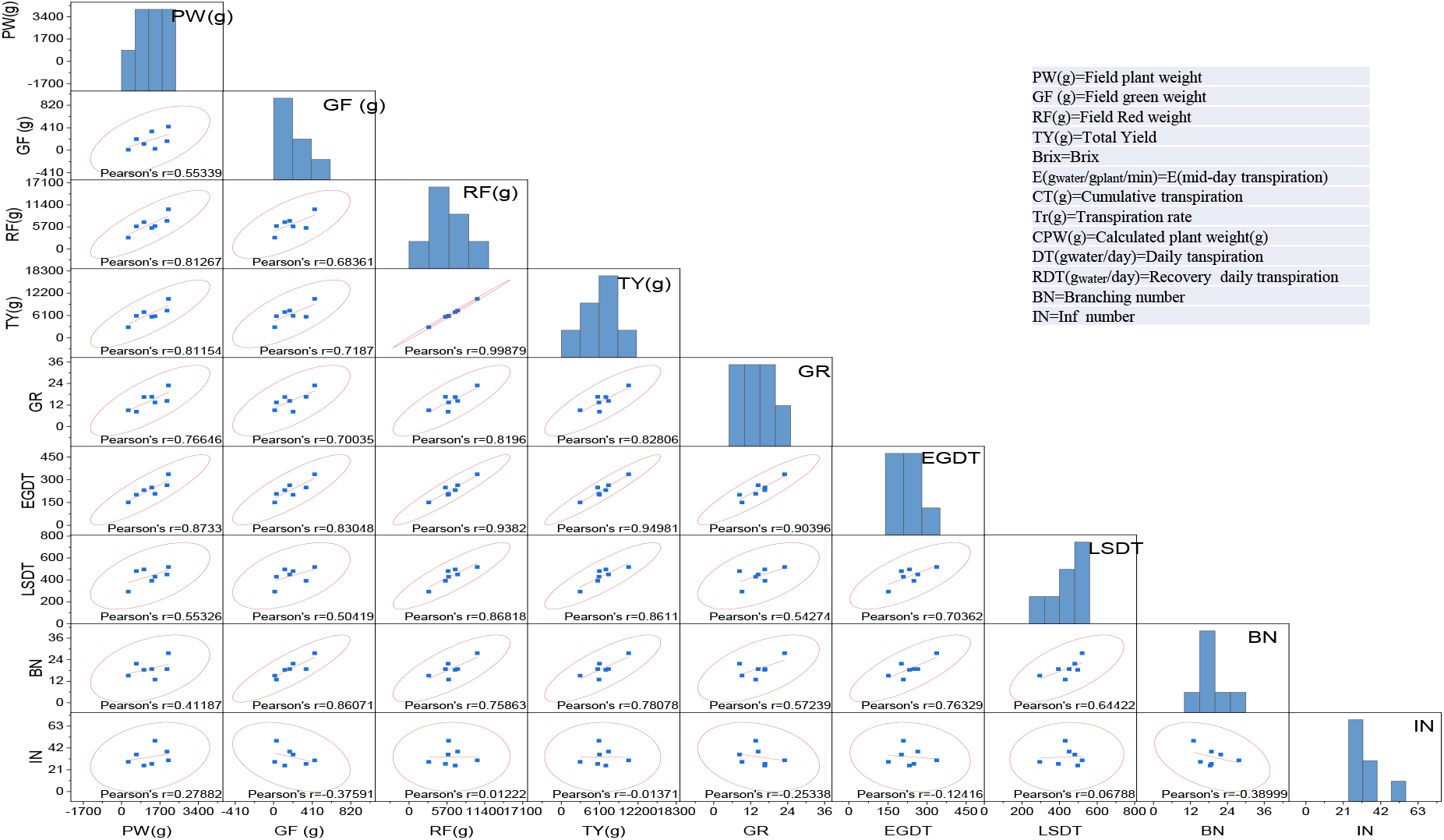
Scatter plot matrix for traits under well irrigated conditions in 2019. The relationships between fields measured yield and its components and greenhouse-based traits of 7 different young tomato plants under irrigated condition. The windows are Pearson’s correlation coefficients (r) and bivariate scatter plots matrix with density ellipse. The short names are defined as below.

**Supplementary Fig. S6.**
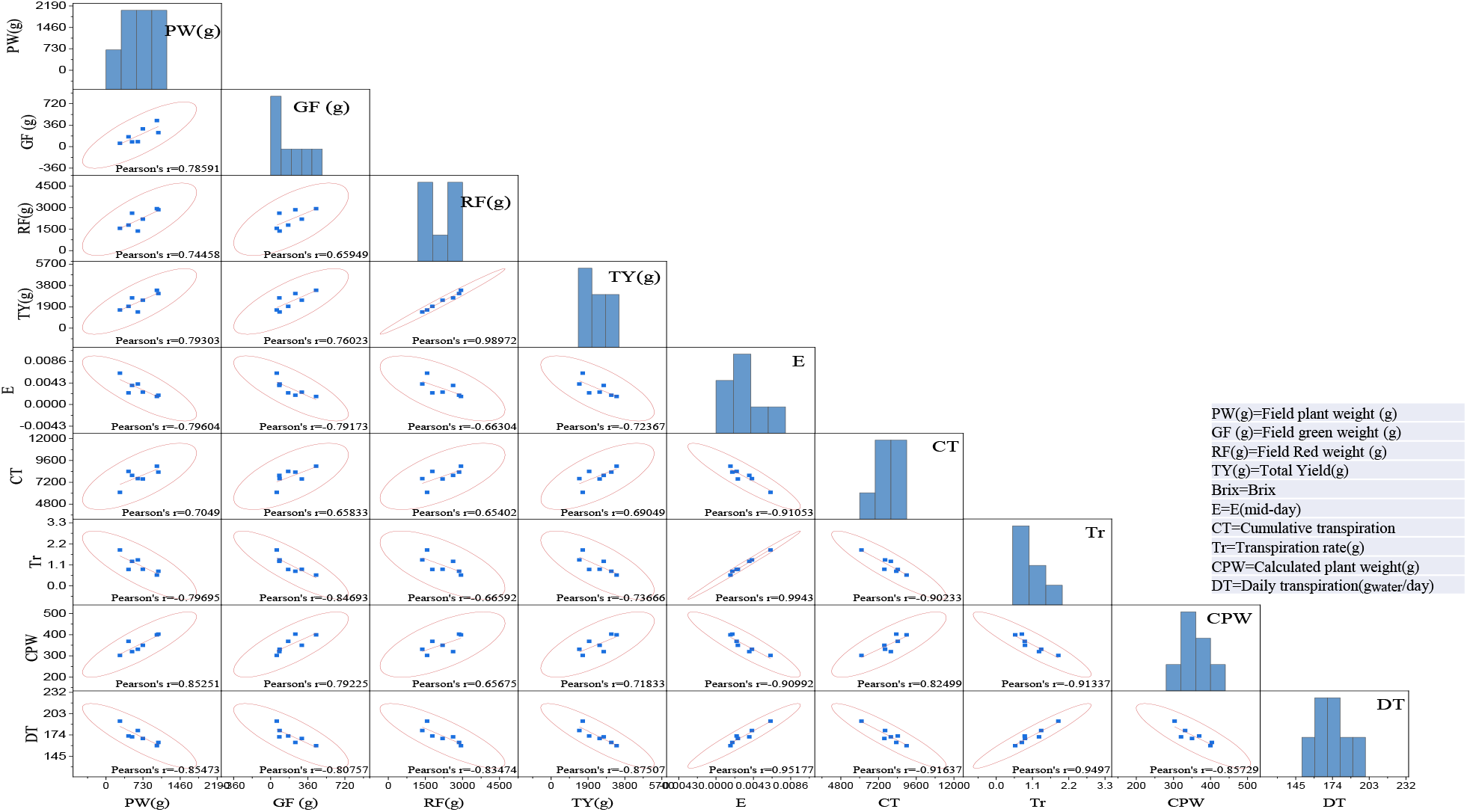
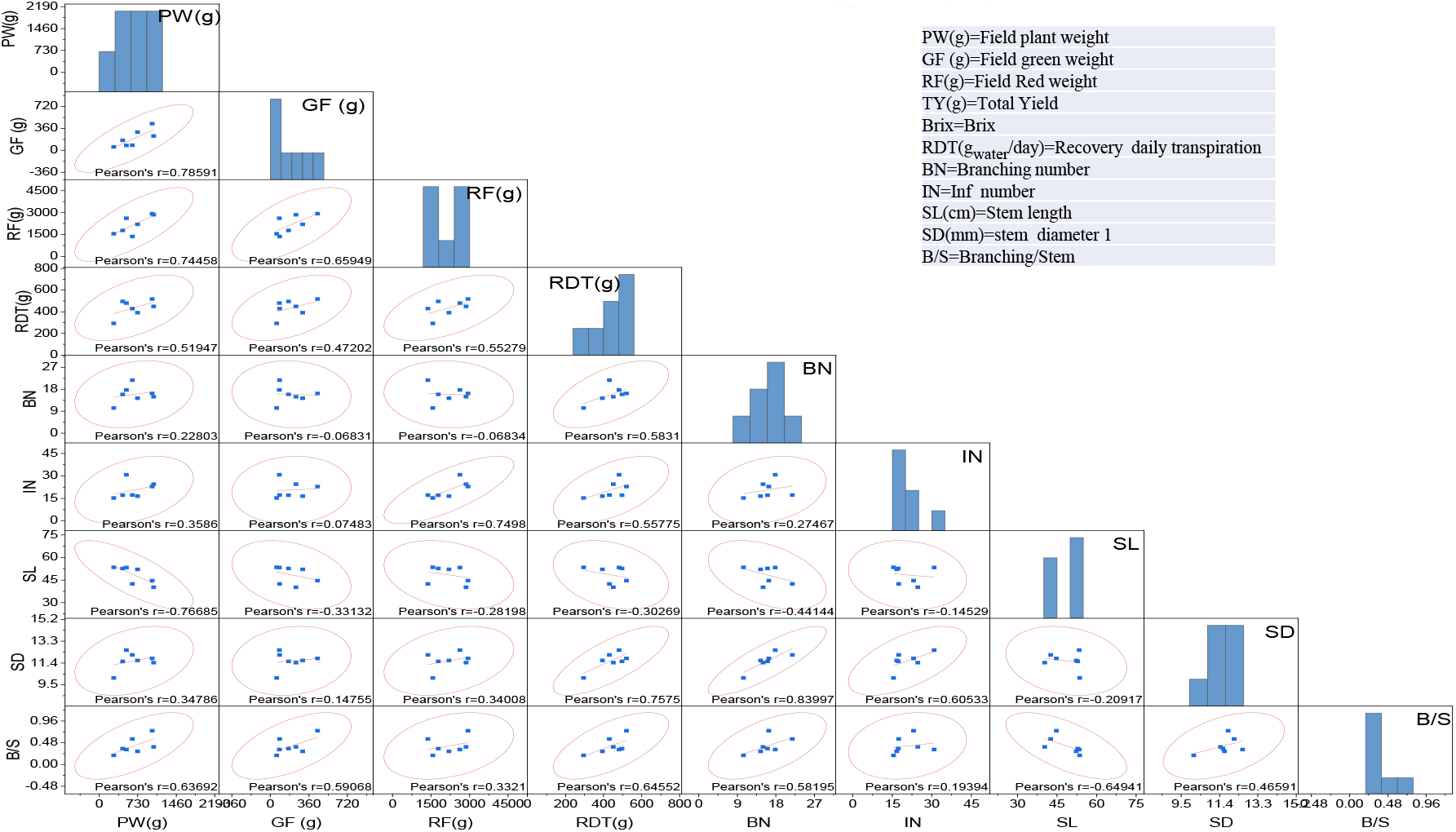
Scatter-plot matrices of correlations among studied physiological yield traits under the water-deficient conditions in 2019. The figure depicts the relationships between field-measured yield and yield components, and the greenhouse-based traits of 7 different young tomato plants under drought condition. The windows show Pearson’s correlation coefficients (*r*) and bivariate scatter-plot matrices with a density ellipse. The short names are defined as below.

## Author contributions

Original research plan conceived by MM, DZ and JH, carried out by SG, AK, and IS. SG, and AK analyzed the data and wrote the MS draft. All authors contributed to the writing, reviewing, and editing of the manuscript, and have read and approved the final version.

## Acknowledgements

The work was funded by the Israel Ministry of Agriculture and Rural Development (Eugene Kandel Knowledge centers) as part of the “Root of the Matter” – The root zone knowledge center for leveraging modern agriculture to MM. This research was also supported by the ISF-NSFC joint research program (grant No. 2436/18) and the United States–Israel Binational Science Foundation (BSF Grant # 2015100) to MM. D.Z. was funded by a TOMRES grant (H2020 #727929). <u>J.H.was funded by I</u>srael Science Foundation, ISF (grant No. 1930/18).

